## Supplemental Figures for "A molecular atlas of plastid and mitochondrial evolution from algae to angiosperms"

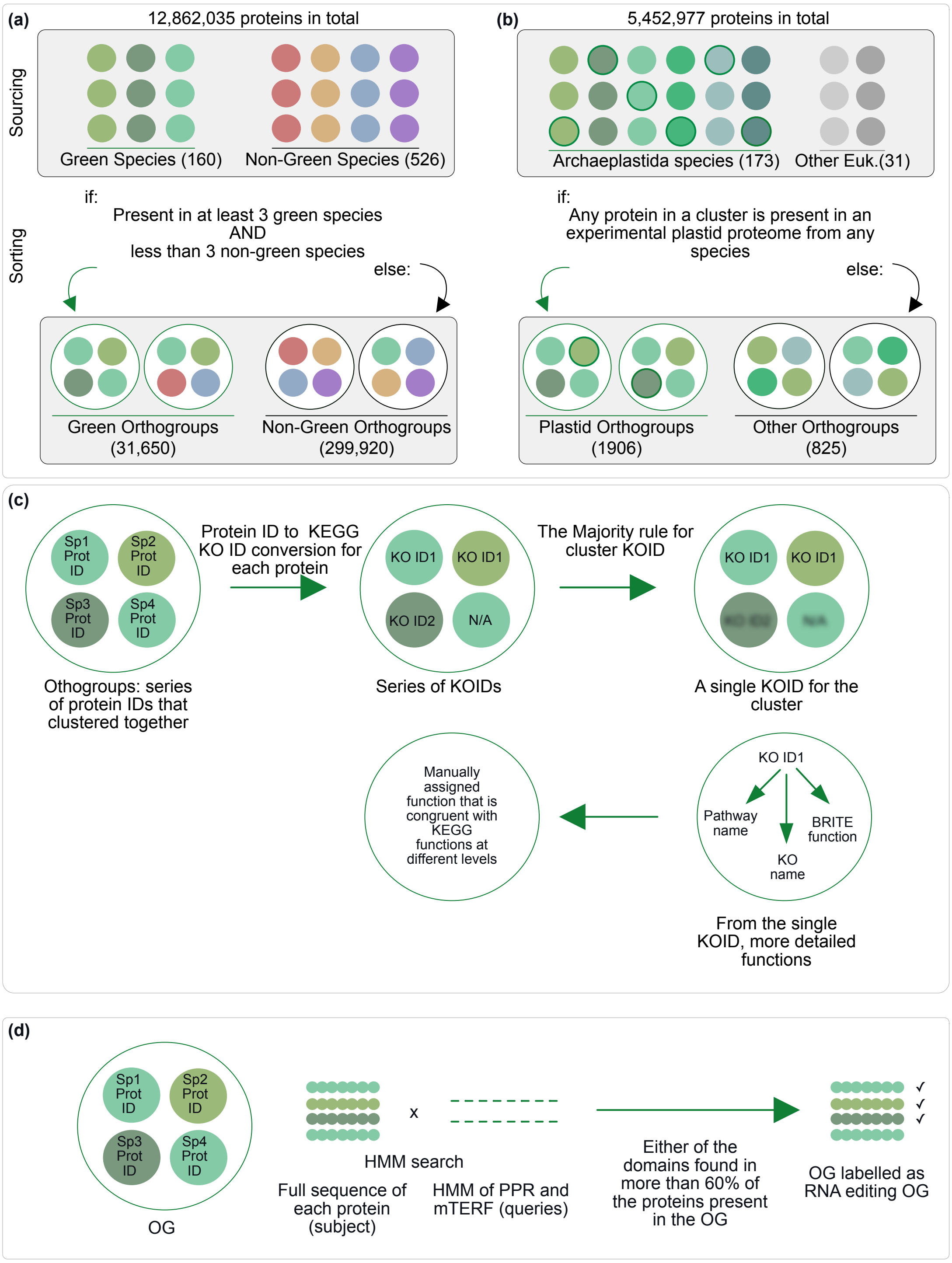


**Fig. S1:** Sorting of (a) the Green OrthoGroups (GOGs) and (b) Plastid OrthoGroups (POGs). In the first step (top boxes), source protein sequences from available species were clustered into protein families. Based on predetermined criteria (between the two boxes), the protein clusters were then separated into the clusters of interest (bottom boxes). Mitochondrial OrthoGroups (MOGs) were sorted the same as **(b)** and based on their presence in any experimental mitochondrial proteome. **(c)** For the functional annotations (for GOGs, POGs and MOGs), KEGG KOID were translated from ProteinID of each protein present in each cluster and from across species. For species outside of the KEGG database (or some proteins within the KEGG database), no KOIDs are available, they are indicated by 'N/A'. The most frequent KOID within a cluster (e.g. KEGG ID 1 in the second circle), was assigned as the KOID for the entire cluster. **(d)** OGs were sorted into PPR or mTERF containing OGs by using the hidden Markov profiles of these domains as a query against all proteins in a given OG. If more than 60% of individual proteins inside an OG contained these domains, we labelled it as RNA editing domain.

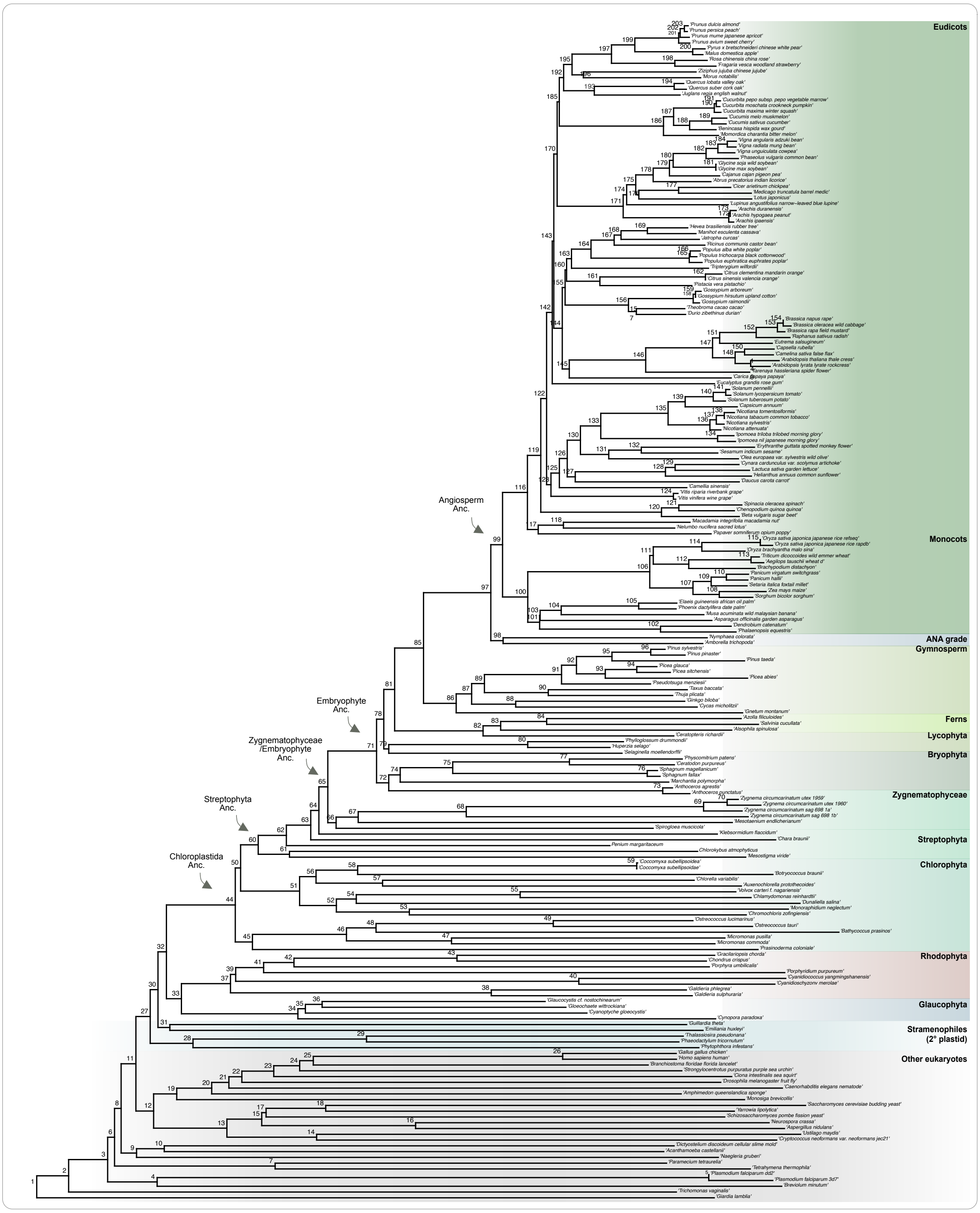


**Fig. S2** Inferred phylogeny of 204 Eukaryotes, with major groups and ancestors of Archaeplastida indicated.

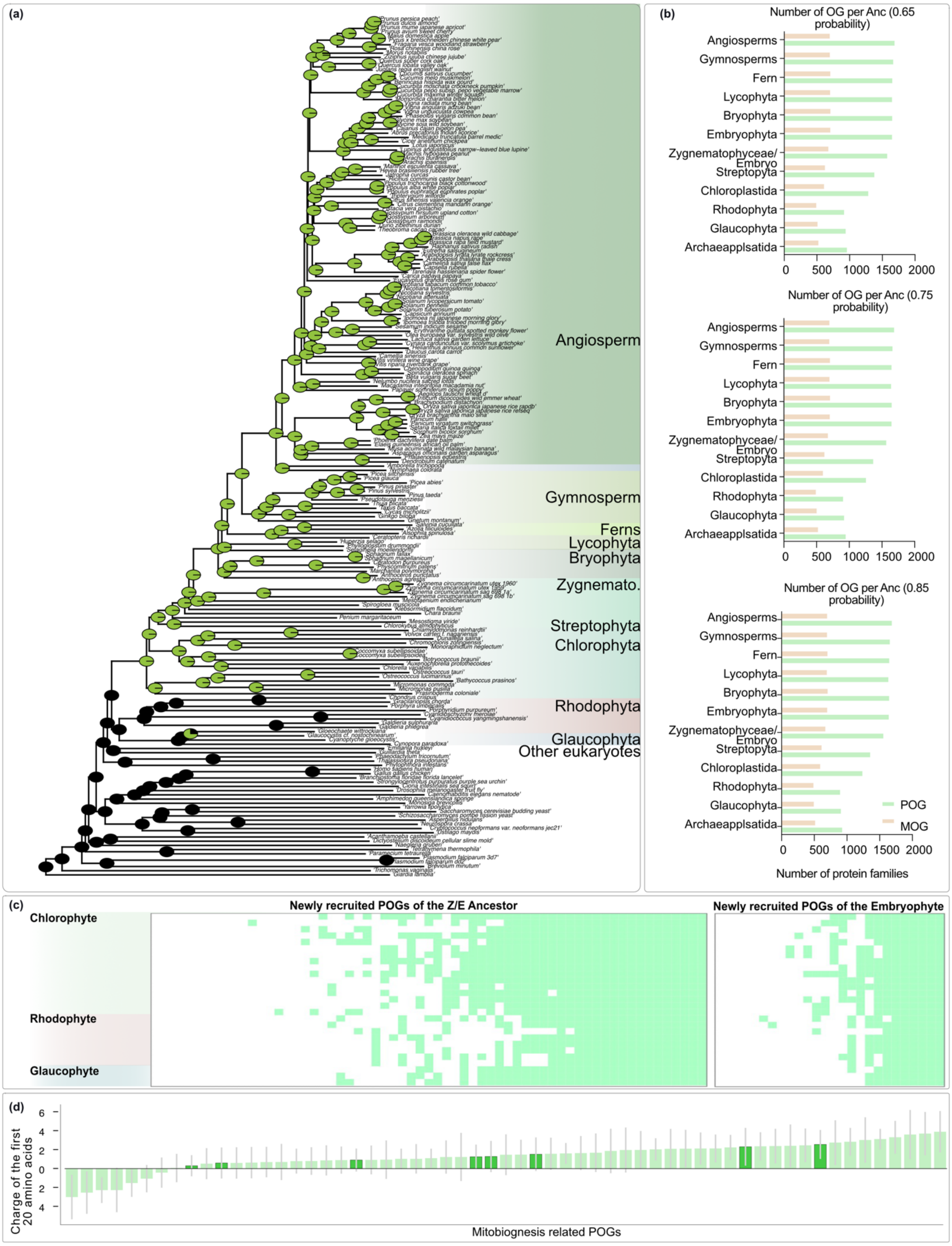


**Fig. S3** Validation of ancestor state reconstruction (ASR) approach on a control protein family of rbcS **(a)**. Number of POGs and MOGs gained by major ancestors, as per probability threshold of inclusion 0.65, 0.75 and 0.85 **(b).** Hidden Markov model-based validation of newly gained POGs of Z/E and Embryophyte ancestor **(c).** Charge of the first 20 amino acids across mitochondrial biogenesis related POGs **(d)**, with mTERF containing POGs present also in MOG shown in a darker shade.

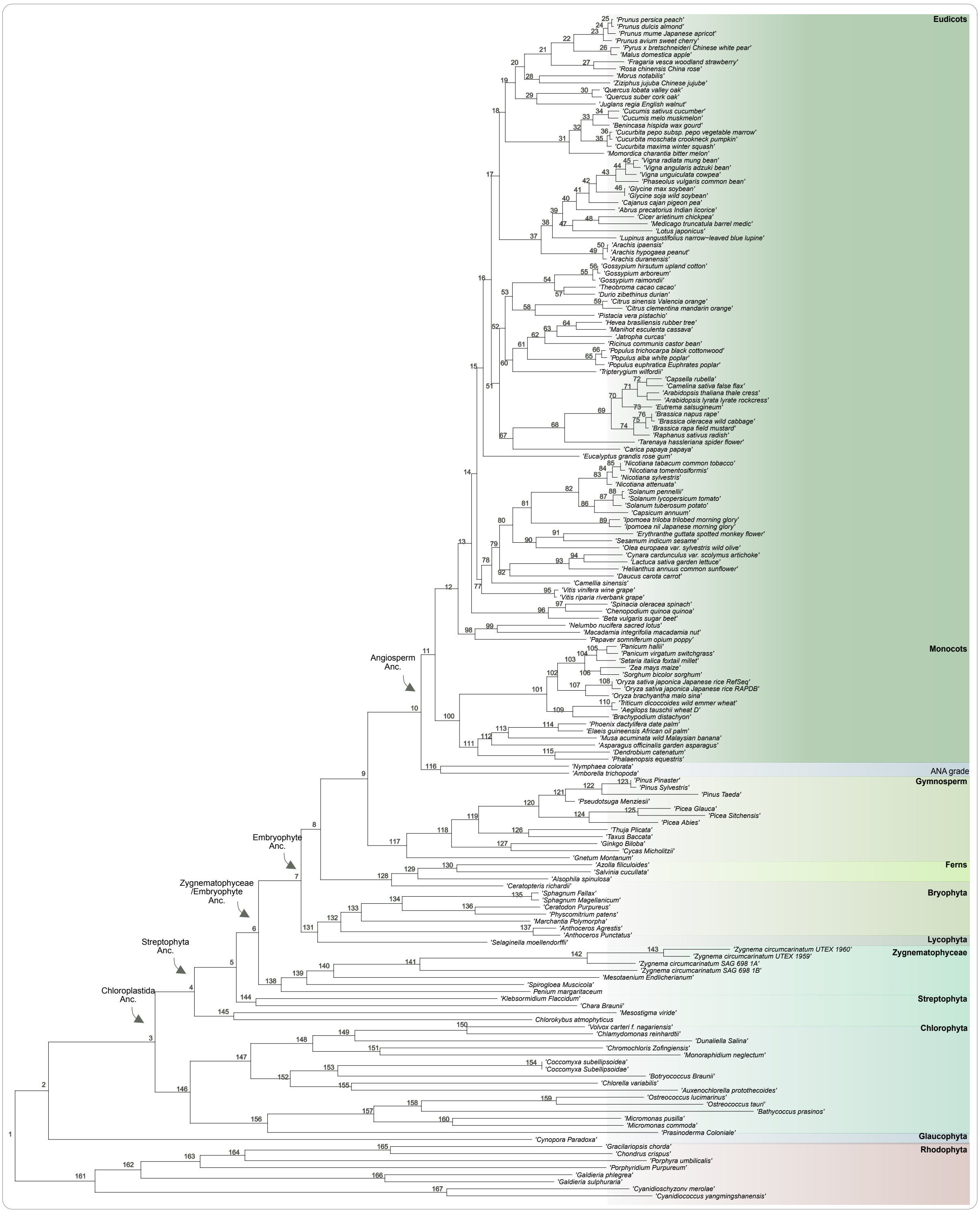

**Fig. S4:** Inferred phylogeny of Archaeplastida (rhodophytes as the sister lineage to all others) with major ancestor nodes indicated with the arrows and major groups highlighted by labels on the right.

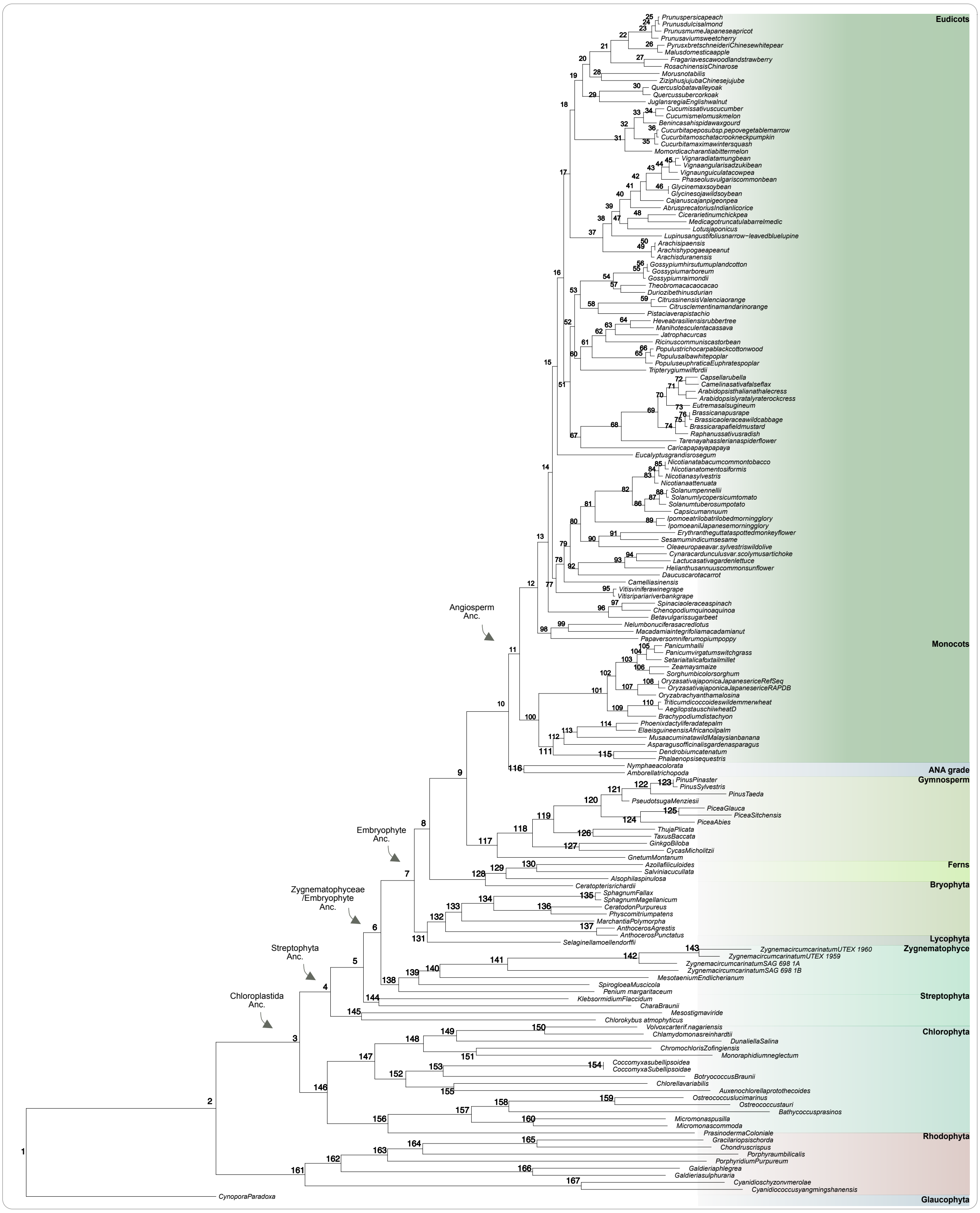

**Fig. S5**: Inferred phylogeny of Archaeplastida (glaucophytes as the sister lineage to all others) with major ancestor nodes indicated with the arrows and major groups highlighted and labelled on the right side.

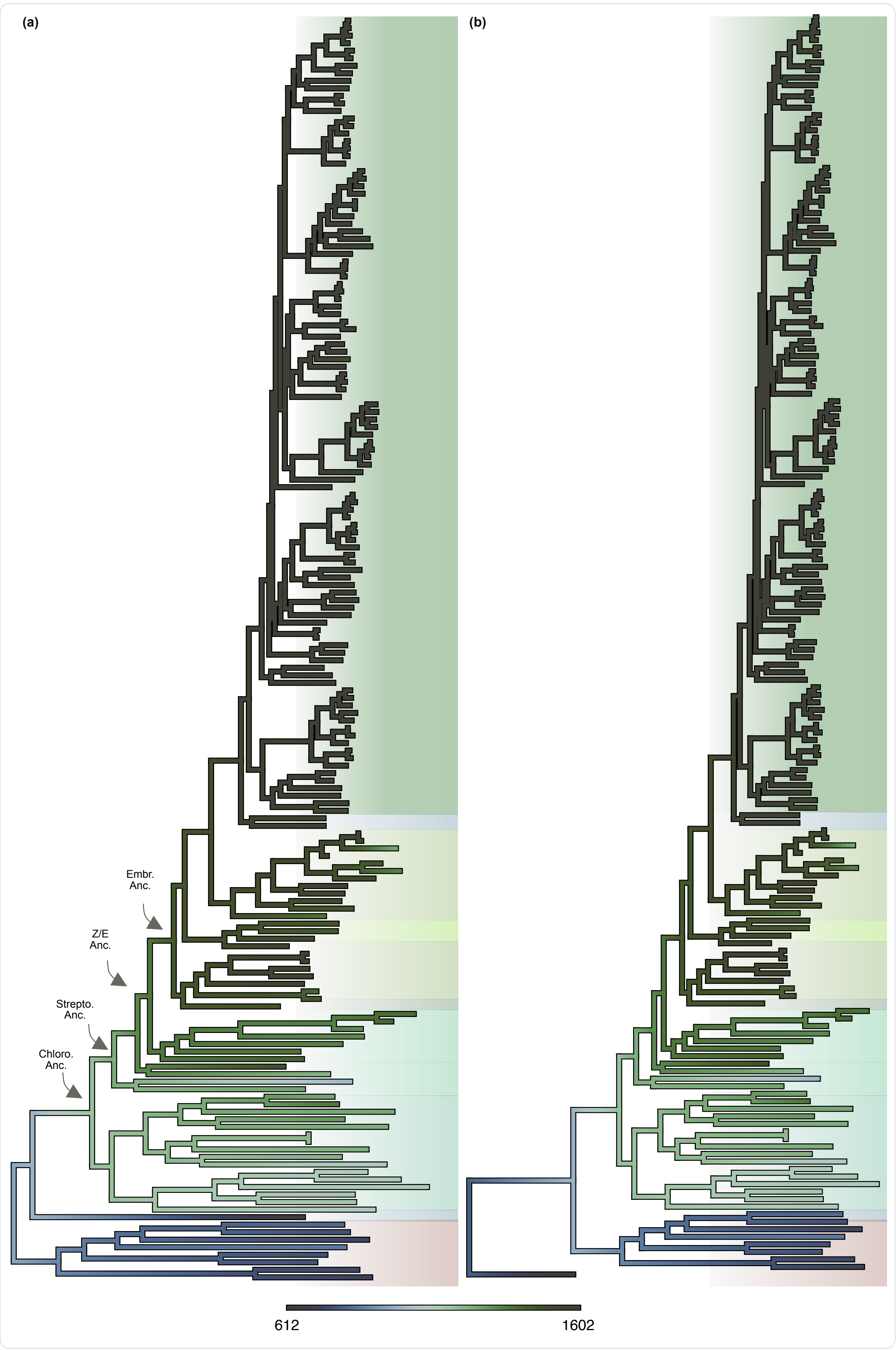


**Fig. S6**: Evolution of plastid orthogroup numbers across the Archaeplastida on a phylogeny with Rhodophyta (a) and Glaucophyta (b) as a basal branch.

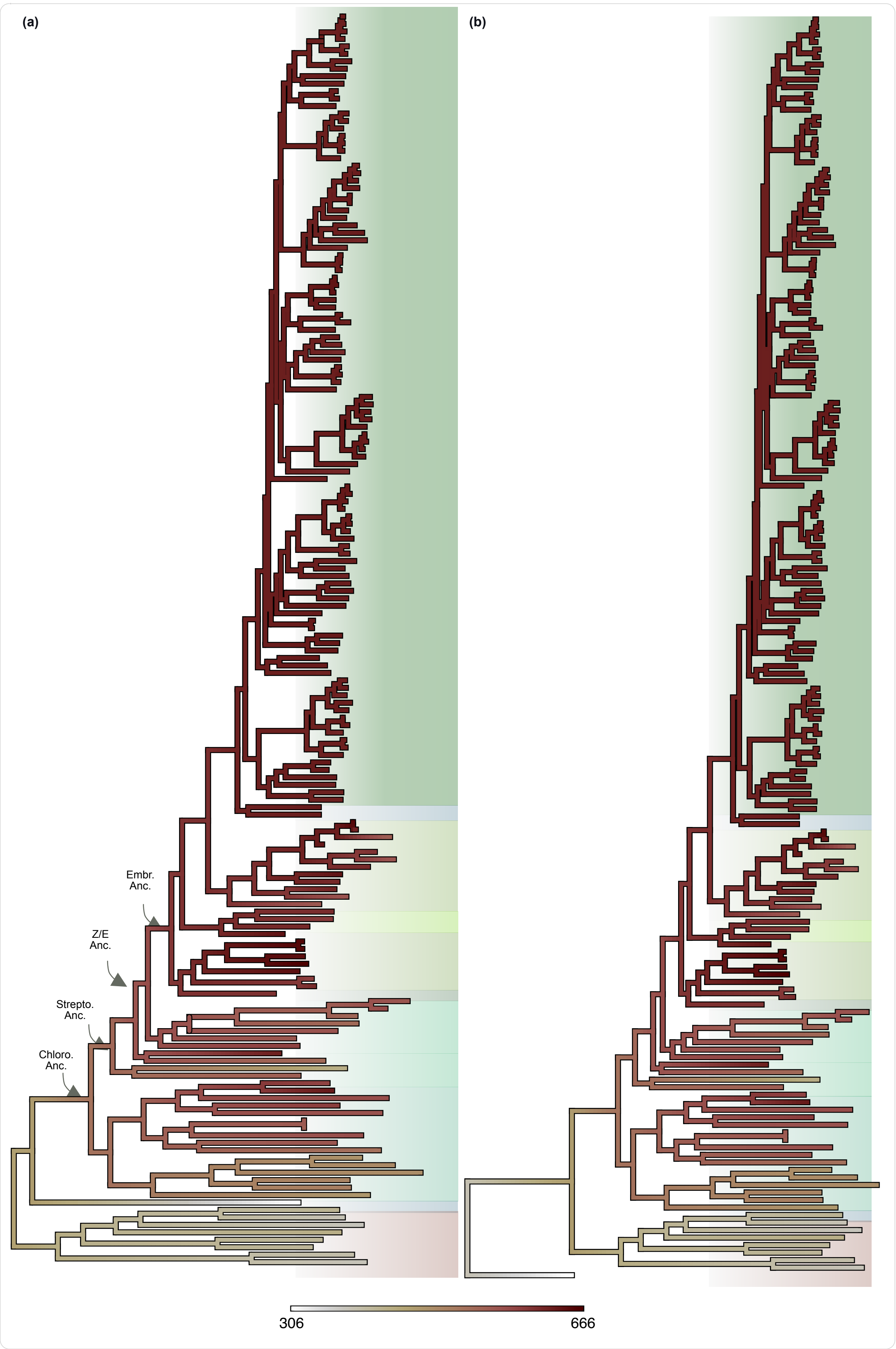


**Fig. S7:** Evolution of Mitochondrial orthogroup numbers across the Archaeplastida on a phylogeny with Rhodophyta (a) and Glaucophyta (b) as a basal branch.

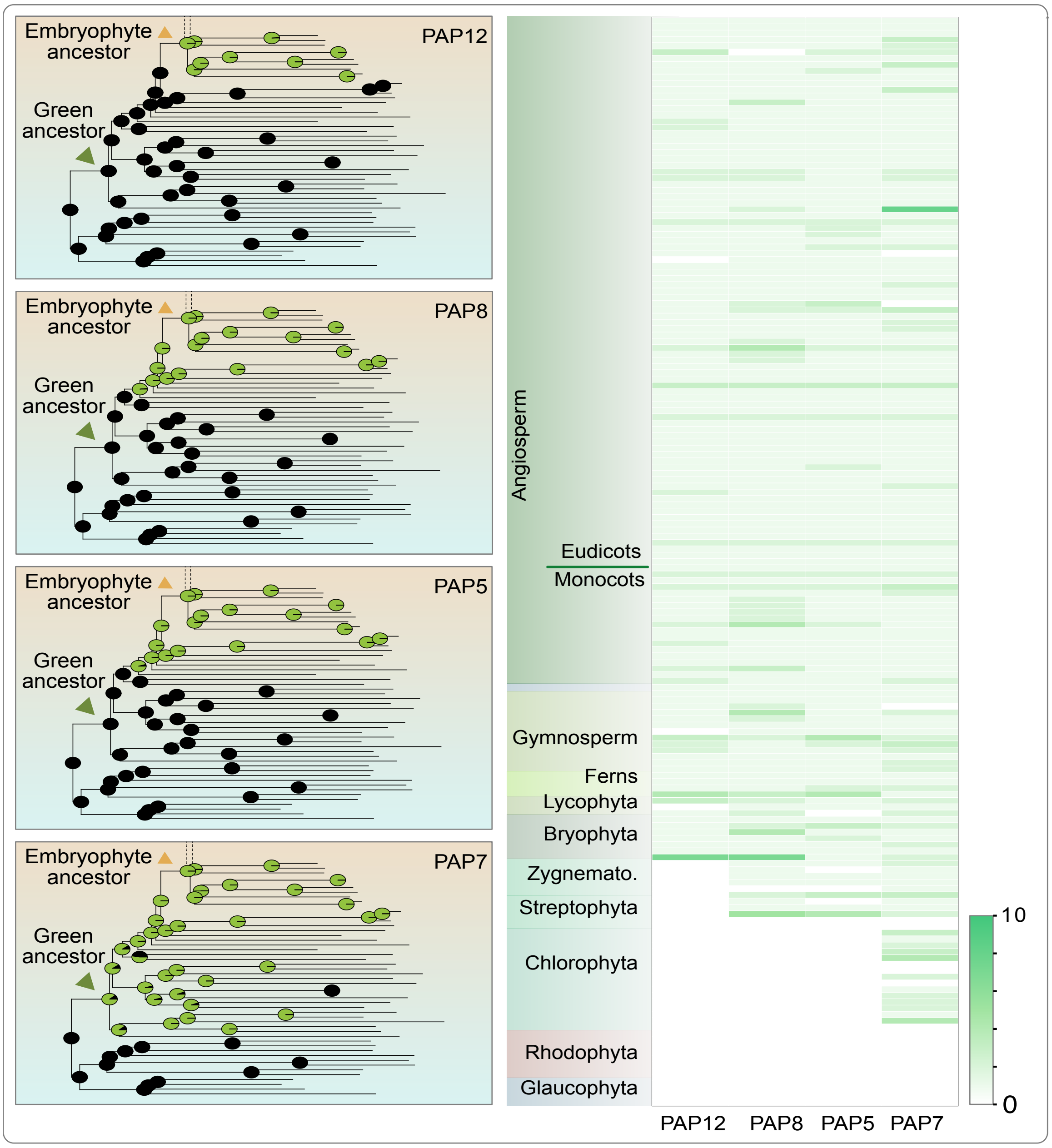


**Fig. S8** Ancestor state reconstruction (ASR) **(a)** and gene copy numbers **(b)** for selected Plastid encoded RNA polymerase interacting proteins (PAPs). The pie charts at each node represent the probability of presence (green) or absence (black) of a protein family in that node.

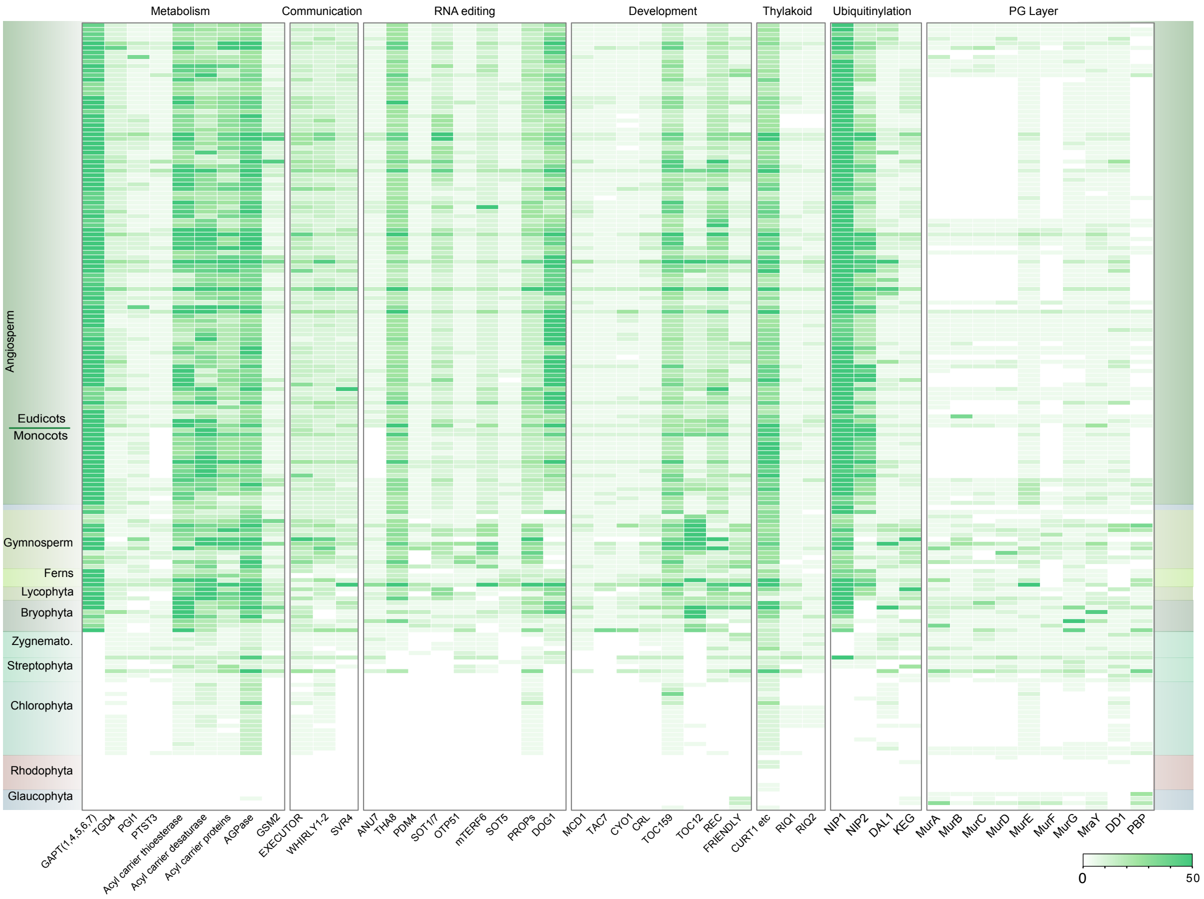


**Fig. S9** Copy number distribution of key proteins recruited around terrestrialization (white indicates absence of a protein, values above 50 were shown as the darkest shade).
